# A three-dimensional carbon – nutrient functional balance model explains the formation of Root Economics Space

**DOI:** 10.64898/2026.04.30.722094

**Authors:** Binbing Zhou, Guangshui Chen

## Abstract

Root carbon (C)–nutrient functional balance underpins root economic strategies, yet previous models neglected the root-length dimension. Here, we develop a three-dimensional C–nutrient functional balance model for first-order roots that explicitly incorporates root length, and validates it using a global root trait dataset. For subtropical woody species, root cortex volume × root length scaled isometrically with the fourth power of stele diameter (slope≈1.0), supporting the model. Herbaceous species from the Qinghai–Tibetan plateau showed a significantly lower slope (0.61), likely due to extreme environmental impacts on transport or metabolism. Woody species preferentially invest in cortical area (thicker roots), supporting mycorrhizal symbiosis, whereas herbaceous species favor root length extension for autonomous soil exploration. By integrating root length, this model provides a novel mechanistic explanation for the formation of both the collaboration and conservation axes within the root economics space, advancing the theoretical framework of root functional strategies.

## INTRODUCTION

Fine roots are the primary organs for acquiring soil nutrients and water, yet their construction, maintenance, and symbiotic associations demand a continuous supply of photosynthates, making them a major carbon (C) sink that can account for 10%–60% of annual gross primary production (Chen *et al*., 2019). This interdependence creates two inextricably linked functional balances within each absorptive root: one between C supply (via phloem) and C consumption (by cortical and fungal metabolism), and another between nutrient absorption (largely mediated by the cortex) and nutrient transport (via xylem). The structural design of roots—particularly the relative investment in cortex versus stele—must therefore simultaneously satisfy these dual constraints of C economy and nutrient logistics (Cao *et al*., 2025).

Two leading mechanistic frameworks have been proposed to explain how root anatomy achieves this balance. The nutrient absorption–transportation balance theory posits that to maintain efficient nutrient delivery to the shoot, the increase in nutrient-absorbing cortical thickness must outpace the increase in stele radius, resulting in an allometric scaling where cortex thickness increases faster than stele radius with increasing root diameter (Kong *et al*., 2017). Conversely, the C supply–consumption balance theory argues that a similar allometry is necessary to balance the C supplied by phloem in the stele with the C consumed by the cortex and associated mycorrhizal fungi (Kong *et al*., 2021; Colombi *et al*., 2022). Both theories are grounded in the physical laws governing fluid flow—notably the Hagen–Poiseuille law (Lambers & Oliveira, 2019), whereby transport rate in conduits scales with the fourth power of vessel radius—and the principle of functional balance between the stele and the cortex.

However, a critical limitation persists: these frameworks, and nearly all empirical tests of them, are confined to a two-dimensional cross-sectional perspective. Roots are inherently three-dimensional cylindrical structures. The longitudinal dimension—root length (L)—has been largely overlooked, despite its dual physiological impacts. Increasing root length expands the total cortical volume (Vc), enhancing potential space for mycorrhizal colonization and resource acquisition (Zheng *et al*., 2024), but it also increases the total metabolic C demand of cortical tissues. Simultaneously, according to the same Hagen–Poiseuille law, increased length raises hydraulic resistance within the stele, potentially reducing transport efficiency (Chen *et al*., 2012). The neglect of this third dimension may explain why the allometric relationship between cortex and stele exhibits substantial variation or is sometimes absent in observational studies (Wang *et al*., 2019; Wang *et al*., 2025c; Zheng *et al*., 2024; Zhou *et al*., 2022). Therefore, a comprehensive functional balance model for absorptive roots must explicitly incorporate the root-length dimension to bridge the gap from microscopic anatomy to macroscopic root system construction strategies.

Under this three-dimensional constraint, where cortical volume that a given stele transport capacity can support is finite, plants face a strategic trade-off: they can invest in increasing cortex cross-sectional area (CCSA) (leading to thicker roots) or in extending root length (leading to longer, thinner roots) (Ma *et al*., 2018). This fundamental trade-off is evident across plant life forms (Eissenstat *et al*., 2000; Meyers *et al*., 2025; Poot & Lambers, 2003; Wei *et al*., 2024).

Woody perennial species often adopt a “thick-and-short” strategy, investing in robust cortical areas to support persistent mycorrhizal symbiosis and durable transport pathways. In contrast, herbaceous species frequently employ a “thin-and-long” strategy, prioritizing rapid soil exploration via extensive root length (Bergmann *et al*., 2020; Wang *et al*., 2025a). Yet, a unified mechanistic model that explains how C and nutrient balance simultaneously shape this cortical area vs. root length trade-off is still lacking.

The three-dimensional constraint is also central to the contemporary paradigm of root functional ecology—the Root Economics Space (RES). The RES organizes global root trait variation along two principal axes: a ‘collaboration gradient’ (typically captured by the negative correlation between specific root length, SRL, and root diameter, RD), reflecting a spectrum from do-it-yourself soil exploration to outsourcing via mycorrhizal fungi; and a ‘conservation gradient’ (captured by the trade-off between root tissue density, RTD, and root nitrogen concentration, RN), reflecting a fast-slow spectrum of resource use (Bergmann *et al*., 2020). While the RES provides a powerful descriptive framework, the mechanistic origins of these two axes, particularly how they arise from the fundamental biophysical and physiological constraints outlined above, remain debated and inadequately explained by existing two-dimensional models (Kong *et al*., 2019; Kou *et al*., 2026). Therefore, a three-dimensional balance model that integrates root length holds the potential to offer new opportunities for elucidating the formation mechanisms of the RES, paving the way for a unified explanation of both the collaboration and conservation axes.

This study focuses on first-order roots, which exhibit high functional homogeneity (Comas & Eissenstat, 2009; Craine *et al*., 2002; Pregitzer *et al*., 2002) and are the most metabolically active and absorptive root class (Gaudinski *et al*., 2010; Huo & Cheng, 2019). Our objectives are threefold: (1) to develop a three-dimensional C–nutrient functional balance model for first-order roots that explicitly incorporates root length, and validate this model using a global dataset of first-order root anatomical and morphological traits; (2) to examine differences in trade-off strategies between woody and herbaceous plants in their investment in CCSA versus root length; and (3) to discuss how this refined model provides a unified mechanistic foundation for the two principal axes of RES—the collaboration axis and the conservation axis.

## MATERIALS AND METHODS

### Theoretical derivation of a three-dimensional C–nutrient functional balance model for first-order roots

We focused on first-order roots and assumed them to be homogeneous cylinders. By incorporating the root length dimension into the classical two-dimensional C–nutrient balance framework (Fig. 1), we examined how root length influences cortical C consumption or nutrient uptake and stele transport rates, and proposed two alternative hypothetical models.

**Fig. 1.**
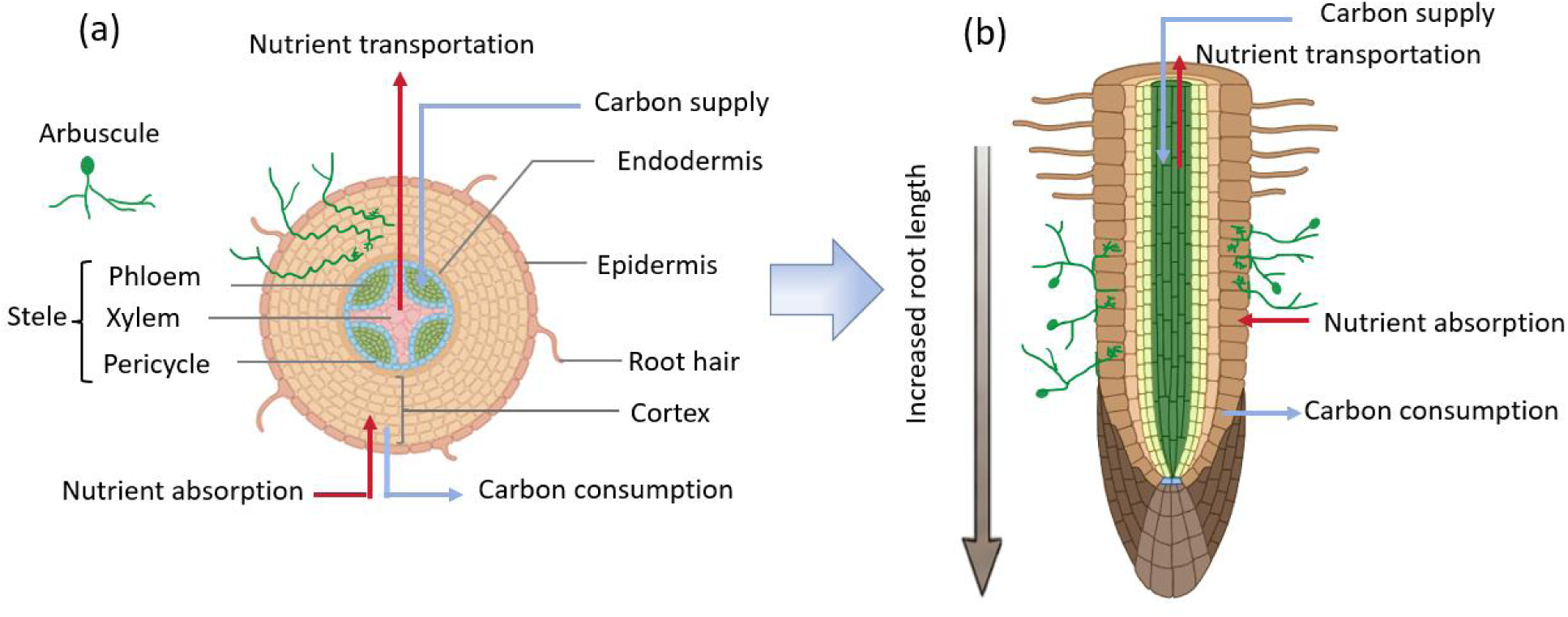
Conceptual diagram of a three-dimensional carbon-nutrient functional balance model, extending from a two-dimensional cross-section (a) to a three-dimensional cylindrical structure (b). This figure was created using BioRender.com under a paid academic license (https://www.biorender.com/).

Hypothesis 1 (Model 1): Only the effect of root length on root cortical C metabolic consumption or nutrient uptake (Y_c_) is considered, while its influence on stele transport capacity (Y_t_) is assumed to be negligible. Under this assumption, the C-nutrient functional balance model in a three-dimensional root volume is formulated as follows:

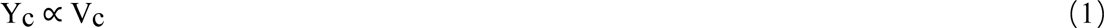

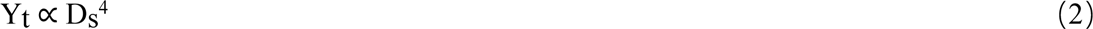

Where V_c_ is the cortical volume and Ds is the stele diameter. According to the Hagen–Poiseuille law (Lambers & Oliveira, 2019), Y_t_ is proportional to the fourth power of Ds.

When transport–uptake (or consumption) is balanced, i.e., Y_c_=Y_t,_ the following relationship is obtained:

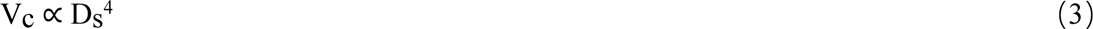

Under this assumption, the V_c_ that a given D_s_ can support is fixed and remains relatively constant regardless of how the root length of this root changes (Fig. 2a).

**Fig. 2.**
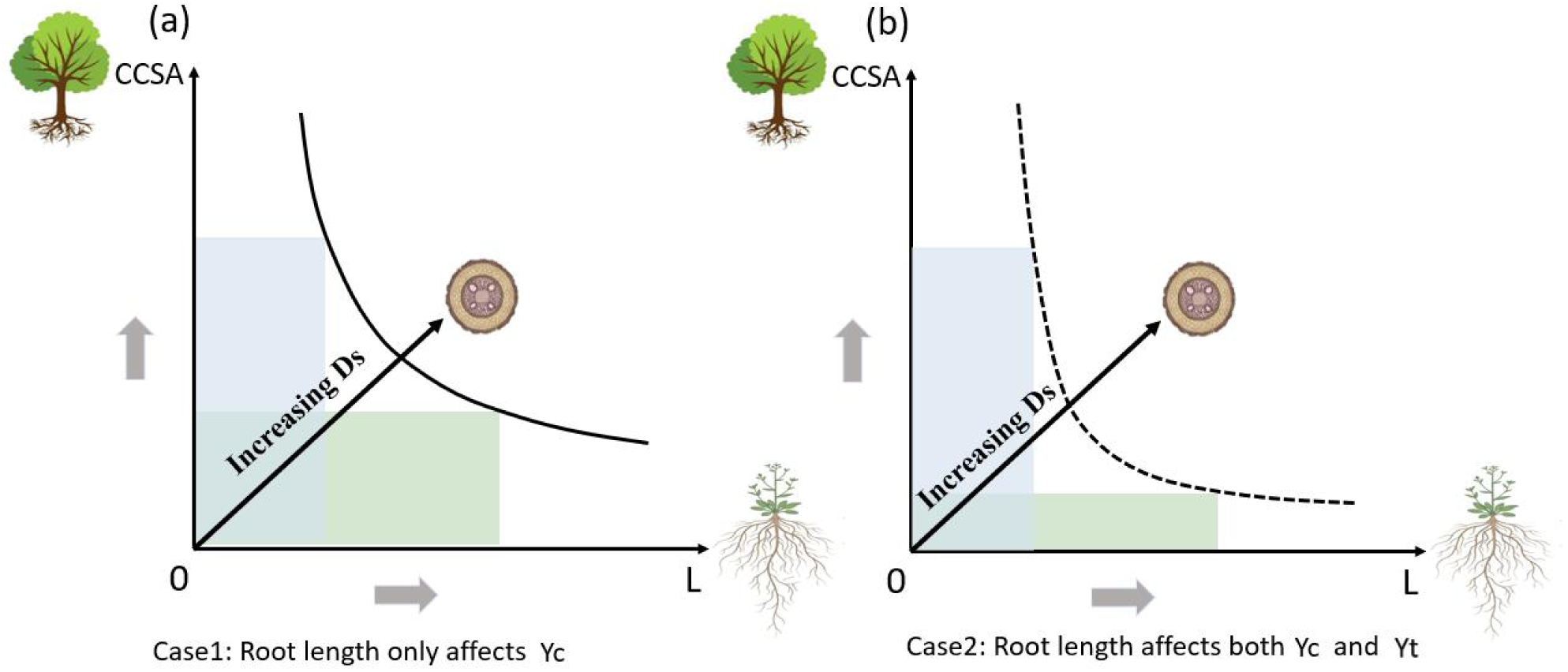
Divergence in cortical and root length investment strategies between woody and herbaceous plants under two hypothetical models. (a) Model 1: Root length does not influence stele transport capacity (Y_t_). Under this case, the cortical carbon consumption or nutrient uptake rate (Y_c_) (indicated by the area of the rectangle formed by the points on the black curve and the coordinate axes X and Y) remain equivalent with increasing root length. (b) Model 2: Increasing root length reduces vascular transport capacity (Y_t_). Under this case, Y_c_ (indicated by the area of the rectangle formed by the points on the black dashed curve and the coordinate axes X and Y) declines with increasing root length. Black arrows indicate the direction of increasing stele diameter. CCSA, cortical cross-section area; L, root length.

Hypothesis 2 (Model 2): The effects of root length on both cortical C metabolic consumption or nutrient uptake (Y_c_) and stele transport capacity (Y_t_) are simultaneously considered. According to the Hagen–Poiseuille law (Lambers & Oliveira, 2019), hydraulic resistance is proportional to conduit length; therefore, stele transport capacity (Y_t_) can be expressed as follows:

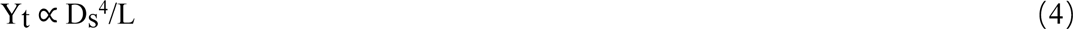

When transport–uptake (or consumption) is balanced, i.e., Y_c_=Y_t_, the following relationship holds:

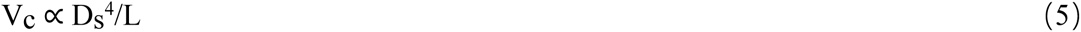

Since V_c_ is calculated as root cross-sectional area multiplied by root length, both terms include the root length variable, which may introduce pseudo-correlation. Therefore, the root length (L) on the right-hand term is transposed to the left-hand term, and the final equation is expressed as follows:

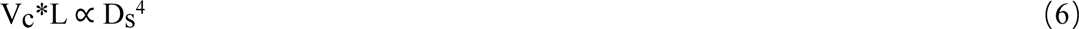

Under this assumption, V_c_ that a given D_s_ can support decreases with increasing root length (Fig. 2b).

Hypothesis 3: For a given D_s_, different plant life forms exhibit variation in investment in root cortex cross-section area and root length due to eco-evolutionary optimization. Using woody and herbaceous plants as an example (Figs. 2, S1), we predict that the allometric scaling exponent of root length versus D_s_ (ɑ) is higher in herbaceous than in woody plants, whereas the scaling exponent of CCSA versus D_s_ (β) is higher in woody than in herbaceous plants. These differences in allometric scaling relationships (Fig. S1) can be expressed as follows:

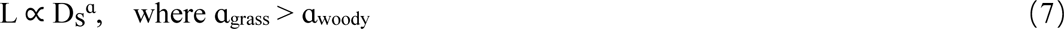

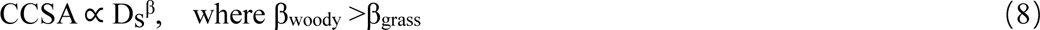

### Data sets

This study systematically compiled anatomical traits (root diameter, stele diameter, cortical thickness) and morphological traits (root length) of plant first-order roots based on three global root trait databases: Fine-Root Ecological Database (FRED), Global Root Trait Database (Groot), and TRY Plant trait Database (TRY). As these databases were last updated in 2019, we conducted a manual search to supplement newly published data from 2020 to 2025 in Web of Science, Google Scholar, Wiley Online Library, and China National Knowledge Infrastructure (CNKI).

For woody plants, we constructed two hierarchical datasets: a regional dataset focusing on subtropical species and a large-scale dataset covering a broader geographic range (see Supplementary Data). For herbaceous plants, which are predominantly distributed in temperate regions, a single integrated large-scale dataset was compiled. It should be noted that there is a severe lack of systematic observation on the length of first-order roots. Apart from the reported data for subtropical woody species (n = 96) (Kong *et al*., 2014), Inner Mongolian herbaceous species (n = 39) (Li *et al*., 2017), and herbaceous species from the Qinghai–Tibetan Plateau (Zheng *et al*., 2024), no additional publicly available data on first-order root length were found. Consequently, when fitting scaling relationships involving root length, only data from these three studies were used. Given the unique geographical environment of the Qinghai–Tibetan Plateau, its data were analyzed separately and were not incorporated into the general herbaceous category. Moreover, the Inner Mongolian herbaceous dataset lacks anatomical measurements; thus, stele diameter was estimated indirectly using an allometric relationship between root diameter and stele diameter derived from the large-scale herbaceous dataset (y = 0.1081 × x¹·¹⁵³⁴, 95% CI of slope: 1.028–1.294, R² = 0.414, P < 0.001, where y is stele diameter and x is root diameter) (Fig. S2). We acknowledge that this estimation introduces uncertainty; however, given the extreme scarcity of first-order root length data, it provides a feasible approach for linking morphological and anatomical traits in a preliminary analysis. In addition, based on the large-scale woody and herbaceous datasets, we separately examined the allometric scaling relationships between root diameter and stele area, as well as between root diameter and cortex area, to further validate differences in resource investment strategies among plant life forms (Figs. S3 and S4).

For studies that reported results only in graphical form, trait data were digitized using GetData Graph Digitizer. For cases where only cross-sectional area was provided, root radius, stele radius, and cortex thickness were back-calculated based on circular geometric relationships. To minimize measurement error, root diameter was consistently defined as the sum of stele diameter and total cortex thickness for all subsequent analyses. When integrating data from multiple sources, duplicate observations for the same species were averaged using arithmetic means. Finally, based on the standardized dataset, we derived additional traits, including cortex cross-section area, stele area, and cortex volume (see Supplementary Data).

### Data analyses

Standardized major axis (SMA) regression was employed to examine scaling relationships among variables, as it provides more robust slope estimation for bivariate relationships (Warton *et al*., 2006). Data analysis was performed on R 4.4.1, using the Smart package.

## RESULTS

### Validation of the three-dimension C-nutrient functional balance model

The results of the standardized major axis (SMA) analysis are presented in Table 1. When considering only the effect of root length on cortical C consumption or nutrient uptake (Hypothesis 1), the regression slope of cortex volume (V_c_) against the fourth power of stele diameter (D_s_⁴) was 0.851 (95% CI: 0.785, 0.922; R² = 0.846, P < 0.001) for subtropical woody species, and 0.452 (95% CI: 0.370, 0.552; R² = 0.341, P < 0.001) for herbaceous species from the Tibetan Plateau. Both slopes were significantly lower than the predicted value of 1, indicating a clear deviation from the expectations of Model 1 (Fig. 3a).

**Fig. 3.**
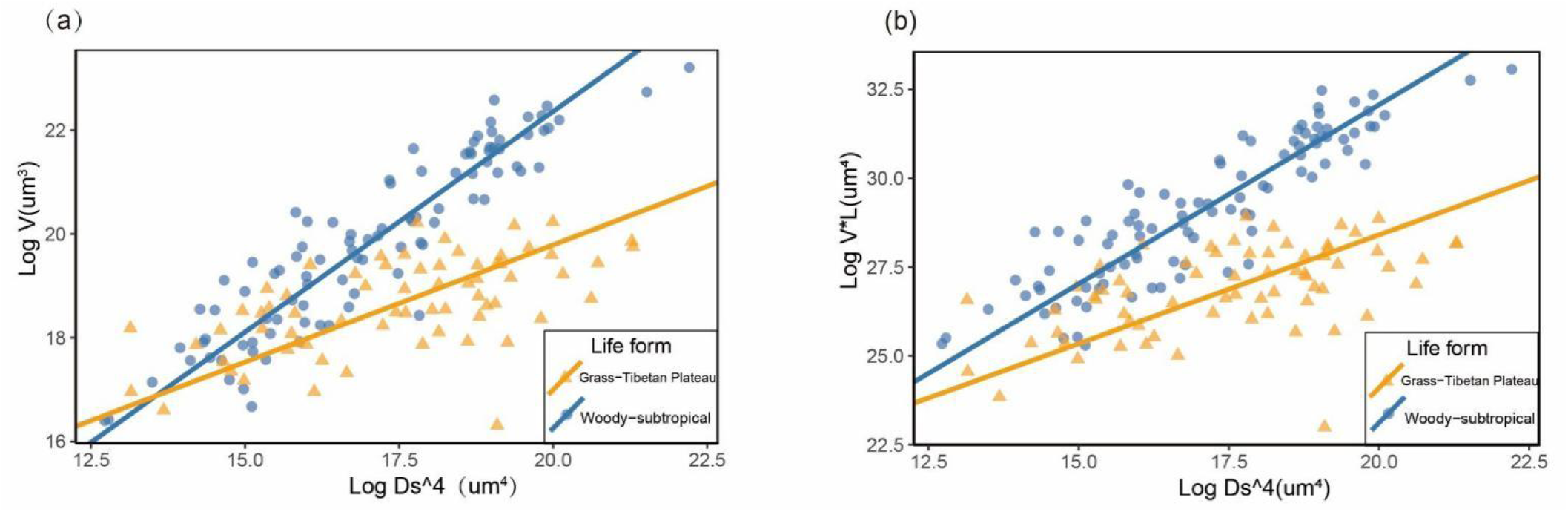
Allometric scaling relationships between root carbon metabolic consumption or nutrient absorptive capacity (represented by cortical volume, V_c_) and stele transport capacity (D_s_^4^) in the first-order roots across different data sources, presented under two scenarios: (a) considering only the effect of root length on cortical carbon consumption or nutrient absorption rate, and (b) considering the dual effects of root length on both transport capacity of root stele and cortical carbon consumption or nutrient absorption rate. Analyses were conducted using standardized major axis (SMA) regression. Subtropical woody samples (n=96) are indicated by blue circles, and Qinghai-Tibetan Plateau herbaceous samples (n=67) are indicated by yellow triangles. Fitting results are presented in Table 1.

**Table 1.** Standardized major axis (SMA) regression results for allometric scaling relationships among first-order root traits.

| Type | Life form | Slope | LowCI | UppCI | Intercept | LowCI | UppCI | $R^2$ | $P$ |
| --- | --- | --- | --- | --- | --- | --- | --- | --- | --- |
| Vc & Ds<sup>4</sup> | Woody-subtropical | 0.8508 | 0.785 | 0.922 | 2.322 | 1.812 | 2.831 | 0.846 | < 0.001 |
|  | Grass-Tibetan Plateau | 0.452 | 0.370 | 0.552 | 4.669 | 3.974 | 5.364 | 0.341 | < 0.001 |
| Vc*L & Ds<sup>4</sup> | Woody-subtropical | 1.008 | 0.919 | 1.105 | 5.173 | 4.478 | 5.869 | 0.796 | < 0.001 |
|  | Grass-Tibetan Plateau | 0.6121 | 0.496 | 0.756 | 7.019 | 6.024 | 8.015 | 0.264 | < 0.001 |
| L & Ds | Woody-subtropical | 0.823 | 0.696 | 0.974 | 2.489 | 2.229 | 2.749 | 0.320 | < 0.001 |
|  | Grass-Inner Mongolia | 1.856 | 1.561 | 2.207 | 0.078 | 0.500 | 0.656 | 0.728 | < 0.001 |
|  | Grass-Tibetan Plateau | 0.740 | 0.584 | 0.937 | 2.162 | 1.825 | 2.500 | 0.075 | < 0.001 |
| CCSA & Ds | Woody-subtropical | 2.877 | 2.664 | 3.107 | -0.717 | 1.130 | 0.303 | 0.859 | < 0.001 |
|  | Woody-global | 2.385 | 2.222 | 2.560 | 0.080 | 0.248 | 0.400 | 0.760 | < 0.001 |
|  | Grass-global | 1.807 | 1.598 | 2.043 | 1.400 | 1.027 | 1.773 | 0.335 | < 0.001 |
|  | Grass-Tibetan Plateau | 1.279 | 1.064 | 1.536 | 2.107 | 1.656 | 2.558 | 0.445 | < 0.001 |

Vc: root cortical volume; L: root length; Ds: stele diameter; CCSA: cortical cross-sectional area.

When simultaneously accounting for the effects of root length on both transport capacity and C consumption or nutrient uptake (Hypothesis 2), the allometric scaling slope between Vc × L and Ds⁴ was 1.008 (95% CI: 0.919–1.105; R² = 0.796, P < 0.001) for subtropical woody species, which is consistent with the predicted value of 1.0 and supports the three-dimensional functional balance model. In contrast, the fitted slope for herbaceous species from the Tibetan Plateau was significantly lower, at 0.612 (95% CI: 0.496–0.756; R² = 0.264, P < 0.001), still deviating from the predicted value of 1.0 (Fig. 3b).

Furthermore, our results indicate that, regardless of whether variation in stele transport capacity with root length is considered, subtropical woody species consistently exhibit higher allometric exponents than herbaceous species from the Tibetan Plateau (Fig. 3; Table 1).

### Divergence in root structural investment among plant life forms

In terms of investment in root length (Fig. 4a), the fitted slope between first-order root length (L) and Ds for herbaceous species from Inner Mongolia was significantly higher than that of subtropical woody species, indicating that herbaceous plants tend to invest more resources to root elongation and rely more on their own uptake capacity for nutrient acquisition. In contrast, the allometric scaling exponent for herbaceous species from the Tibetan Plateau was comparable to that of subtropical woody species, which may reflect a special case under extreme environmental conditions. Regarding cortical investment (Fig. 4b), the scaling slopes between CCSA and D_s_ were consistently higher in woody species than in herbaceous species, suggesting that woody plants preferentially invest in thicker cortex tissue to facilitate mycorrhizal colonization and enhance nutrient acquisition via symbiotic associations. Furthermore, with increasing root diameter, there was a faster increase in CCSA than in stele cross-sectional area in woody species (Fig. S3), whereas similar increase rates between them in herbaceous species (Fig. S4), providing additional support for the patterns observed above.

**Fig. 4.**
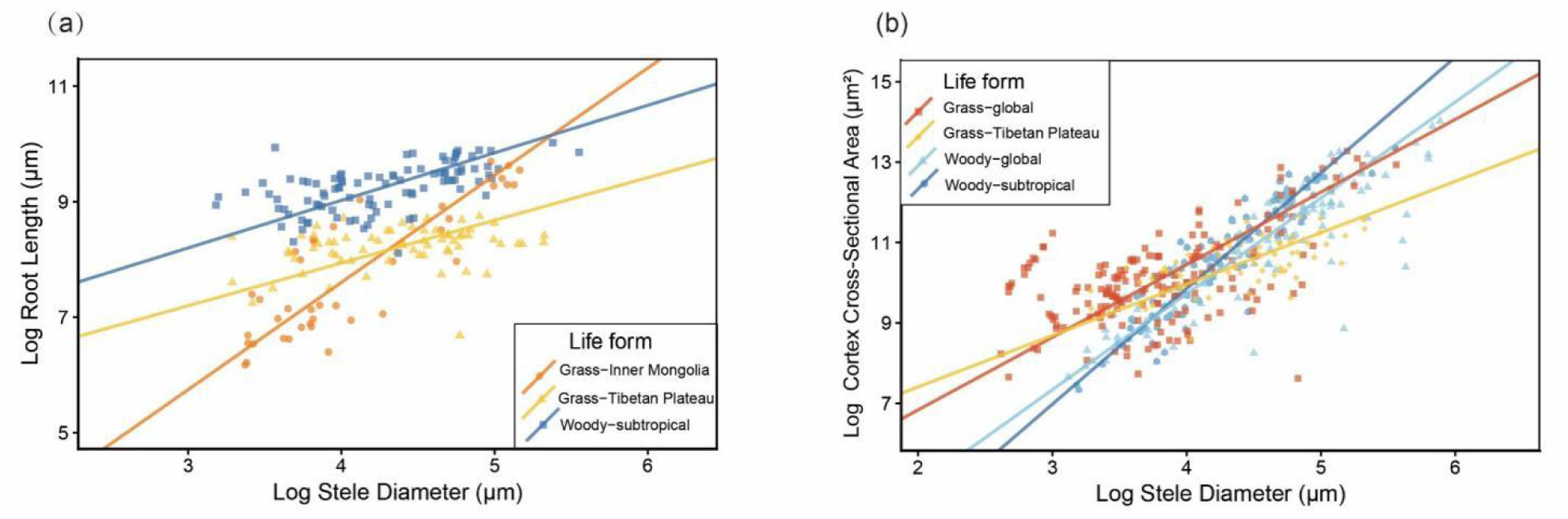
Allometric relationships between (a) root length and stele diameter, and (b) cortex cross-section area and stele diameter for the first-order roots. Subtropical woody samples (n=96) are shown in dark blue, the large-scale woody dataset (n=189) in light blue, Inner Mongolian herbaceous samples (n=39) in orange, Qinghai-Tibetan Plateau herbaceous samples (n=67) in bright yellow, and the large-scale herbaceous dataset in red. Fitting results are presented in Table 1.

Comparisons across datasets from different sources revealed that herbaceous species from the Tibetan Plateau consistently exhibited the lowest allometric scaling slopes (Fig. 4a, b; Table 1). In addition, the scaling slope of CCSA vs. D_s_ in subtropical woody species was higher than that derived from the large-scale woody dataset (Fig. 4b; Table 1).

## DISCUSSIONS

### A three-dimensional C–nutrient functional balance model of the first-order roots

This study shows that, after incorporating the root length dimension, the fitted scaling exponent between Vc × L and Ds⁴ for the first-order roots of subtropical woody species is close to the predicted value of 1, supporting the three-dimensional C–nutrient functional balance model (Hypothesis 2). This finding indicates that root length plays a critical role in regulating resource transport rates in first-order roots, thereby refining the traditional view that emphasizes only stele diameter (including conduit diameter and number) as the primary determinant of transport capacity (Zhou *et al*., 2021), while neglecting the constraint imposed by transport distance (Steudle & Peterson, 1998). The resistance to transport associated with root length is likely linked to conducting tissues. Xylem vessels or tracheids and phloem sieve tubes or sieve cells constitute the primary transport pathways (Roth, 1996; Shen *et al*., 2025), through which photosynthates and nutrients must traverse the end walls and pits within these tissues (Hacke *et al*., 2004; Sperry & Hacke, 2004). As root length increases, the number of end walls and pits encountered rises, thereby augmenting the resistance to mass flow (Chen *et al*., 2012), which may ultimately reduce the transport capacity of the roots.

However, the fitted slopes for the relationships Vc*L ∝ Ds⁴ and Vc ∝ Ds⁴ in herbaceous species from the Qinghai–Tibetan Plateau were both significantly lower than 1. Furthermore, the scaling exponents for the relationships between D_s_ and L, as well as between D_s_ and CCSA, were also lower than those observed in other herbaceous datasets (Fig. 4a, b). We argue that this pattern arises because the extreme alpine environment profoundly alters the transport–consumption balance (Zheng *et al*., 2024). First, low temperatures increase sap viscosity and the risk of xylem embolism, thereby reducing the transport efficiency of the stele (Colmer & Greenway, 2011; Kotula *et al*., 2015; Li *et al*., 2019; Weih & Karlsson, 2001). In response, plants may allocate more resources to the stele, resulting in larger stele diameters and the development of wider or more numerous conduits to enhance transport capacity (Preston *et al*., 2006; Sperry *et al*., 2008). Empirical evidence showing that plants in the Qinghai–Tibetan Plateau generally exhibit larger root diameters and lower specific root length supports this hypothesis (Zheng *et al*., 2025). Second, the extreme environment (low temperature, hypoxia) suppresses the metabolic activity of cortical cells, leading to a relative reduction in cortex volume. Regardless of the specific mechanism, these patterns reflect how extreme environmental conditions compel plants to shift their functional balance point for resource transport and consumption, resulting in a reduced slope that deviates from theoretical predictions.

### Divergent root investment strategies among plant life forms

Our results clearly reveal a trade-off in root investment between woody and herbaceous plants, supporting Hypothesis 3. The “short and thick strategy” exhibited by woody plants (characterized by high CCSA investment and low investment in root length) aligns with their perennial life history and reliance on mycorrhizal symbiosis. Comparative studies on mycorrhizal colonization corroborate this perspective. For example, the first-order roots of *Larix gmelinii* in northeastern China exhibit a 21.9% increase in root diameter following colonization, accompanied by decreases of 23.7% in root length and 16.3% in specific root length (Sun, 2007). In maize, inoculation with different arbuscular mycorrhizal fungal strains also led to significant reductions in SRL(Zhang *et al*. 2025). Herbaceous species such as *Allium porrum* and *Andropogon gerardii* also exhibit trends toward increased root thickening and reduced root length following artificial enhancement of mycorrhizal colonization (Berta *et al*., 1990; Jackson & Caldwell, 1993). Although the mycorrhizal colonization effects vary with mycorrhizal type, that is, arbuscular mycorrhizal (AM) fungi tend to promote root thickening whereas ectomycorrhizal (EM) fungi more strongly enhance root branching (Berta *et al*., 1995; Rousseau *et al*., 1994; Yan *et al*., 2022; Ding *et al*., 2023), the core strategy of woody plants is to prioritize the symbiotic space rather than extending root length. At the same time, by reducing investment in root length, woody plants may decrease hydraulic resistance along the transport pathway, thereby potentially achieving higher rates of C and nutrient transport (Fig. 2). This strategy also supports the high C demand and elevated nutrient transport rates required to sustain mycorrhizal symbiosis.

In contrast, herbaceous plants, characterized by short life cycles and intense competition, generally adopt a “do-it-yourself” resource acquisition strategy. They preferentially invest in root length, relying on rapid soil exploration to occupy space and capture resource patches (An *et al*., 2021; Einsmann *et al*., 1999; Wang *et al*., 2025a; Yan *et al*., 2025). This strategy enables herbaceous species to thrive under pronounced environmental fluctuations, such as seasonal drought or cold stress (Ma *et al*., 2018). Consequently, these plants occupy the high specific root length (SRL) and high root nitrogen concentration (RN) end of the root economics space (Wang *et al*., 2025a). To maintain a certain transport efficiency while achieving rapid elongation, herbaceous plants may not invest in cortical thickening as extremely as woody plants relative to the stele (Fig. S3); instead, they exhibit an isometric scaling between cortex and stele dimensions (Fig. S4). As a result, even at comparable root diameters, herbaceous plants maintain a thinner cortex, which likely contributes to their lower mycorrhizal colonization rates relative to woody plants (Ma *et al*., 2018). Furthermore, reduced investment in cortical tissues may also represent an adaptive response to drought stress, as it effectively lowers the resistance to water and nutrient influx into root tissues (Wang *et al*., 2025b) and reduce C costs associated with cortex maintenance (Kong *et al*., 2017, 2021). Thus, the C-nutrient balance in herbaceous species appears to be regulated more through “root elongation” rather than “root thickening” (Fig. 2), enabling herbaceous plants to persist under C- and water-limited conditions.

In addition, this study found that the scaling exponent of the relationship between D_s_ and CCSA in subtropical woody species is higher than that reported for the global woody dataset (Fig. 4b). This discrepancy is primarily attributable to differences in sample representation. Subtropical woody species rely more heavily on mycorrhizal symbiosis for nutrient acquisition (Ma *et al*., 2018), leading to greater investment in cortical tissues. In contrast, the large-scale dataset includes 189 tree species spanning from tropical to boreal regions (see Supporting Information Data) and has a higher proportion of ectomycorrhizal (EM) species (18% vs. 8% in the subtropical dataset). Owing to differences in investment patterns associated with mycorrhizal types (Rousseau *et al*., 1994; Yan *et al*., 2022), the overall fitted scaling exponent is consequently lower for subtropical woody plants. These results further indicate that environmental factors (e.g., climate, soil) and mycorrhizal types play key regulatory roles in shaping root investment strategies in plants.

### Implications of the three-dimensional C–nutrient functional balance model for the formation mechanisms of the Root Economics Space

The two-dimensional RES has become the dominant framework for investigating variation in root form and function (Bergmann *et al*., 2020; Fan *et al*., 2024; Matthus *et al*., 2025). Although the bi-dimensionality of the RES has been partially explained (Kong *et al*., 2019; Zhang *et al*., 2024), the underlying mechanisms driving the collaboration axis and the conservation axis remain insufficiently resolved. The three-dimensional C–nutrient functional balance model developed in this study aims to provide a unified mechanistic insight into the formation of both axes. The model posits that root function is governed by a physical constraint: the balance between the stele-determined transport capacity (Yt) and the cortex-mediated metabolic and absorptive capacity (Yc). Within this overarching constraint, plants differentiate their investment strategies at two distinct levels (organ and cellular level), thereby shaping the two dimensions of the RES.

Traditionally, the “collaboration axis” has been characterized by the negative relationship between root diameter (RD) and specific root length (SRL), interpreted as a trade-off between “do-it-yourself” and “mycorrhizal-dependent” strategies. However, this relationship may largely arise from geometric constraints of cylindrical structures of roots (Zhang *et al*., 2024), rather than reflecting a fundamental functional divergence. For instance, the RD-SRL trade-off persists in higher-order roots where cortex is reduced and mycorrhizal collaboration is irrelevant (Kou *et al*., 2026). We therefore propose that the essence of the collaboration axis is more accurately captured by a trade-off between CCSA and L (or L^2^) (Fig. 5). This directly corresponds to the core constraint of our three-dimensional model: under the maximum transport capacity (Yt ∝ Ds⁴, or Yt ∝ Ds⁴/L) set by stele diameter, the total cortex volume (Vc) that can be supported is finite.

**Fig. 5.**
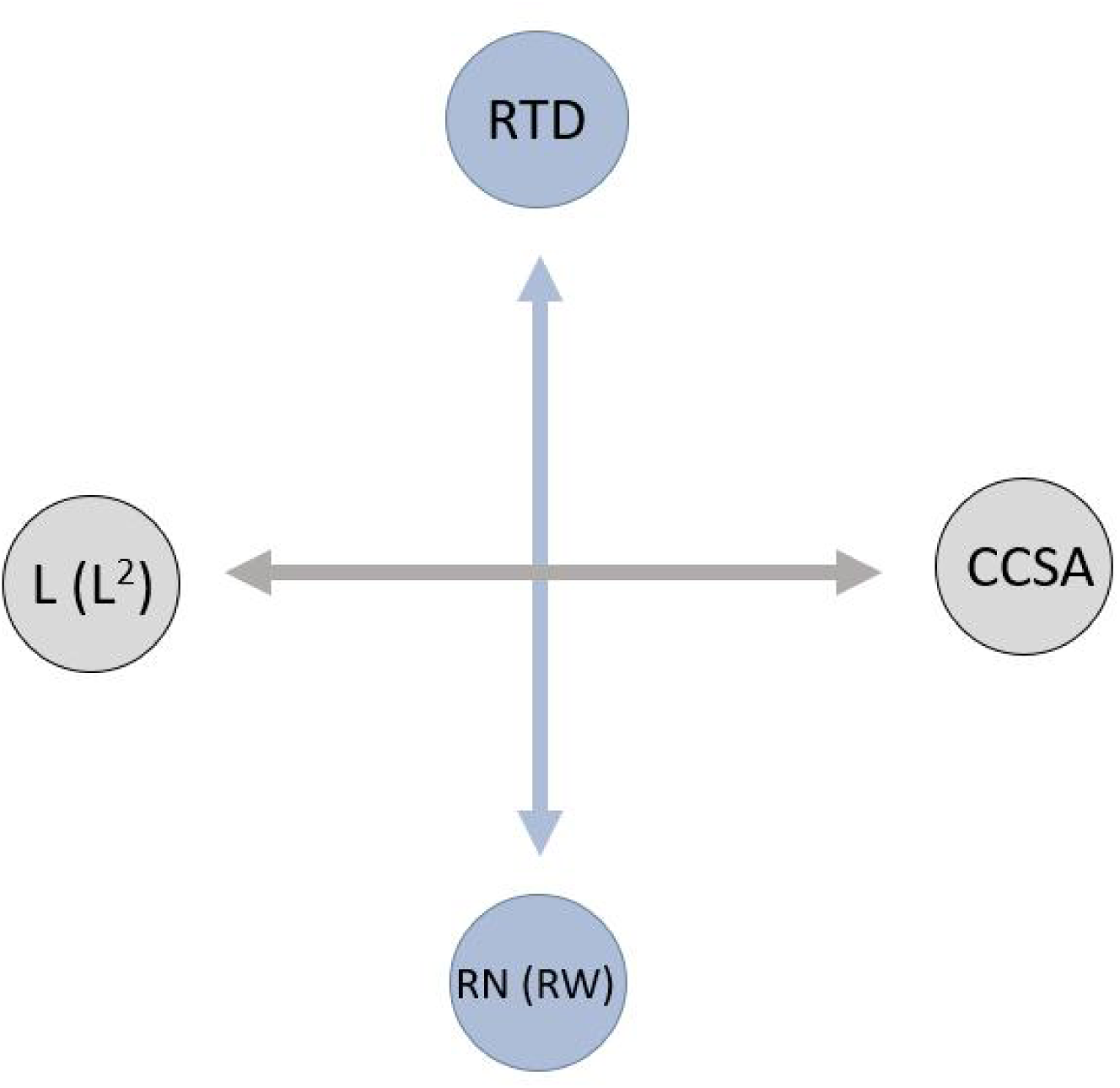
The proposed scheme for the conservation axis and the collaboration axis in root economic space. L, root length; CCSA, cortex cross-sectional area; RTD, root tissue density; RN, root nitrogen concentration; RW, root water concentration. The conservation axis and the collaboration axis are orthogonal to each other.

Plants must therefore trade-off between investing in a thicker cortex (high CCSA) to provide ample space for mycorrhizal fungi (an “outsourcing” strategy), and investing in greater root length (high L) to independently explore a larger soil volume (a “do-it-yourself” strategy) (i.e., Vc=CCSA×L≈ constant). In contrast, RD conflates information from both cortex and stele, while SRL is strongly influenced by RTD (Kong *et al*., 2019), which can obscure or even reverse its relationship with RD (Zheng *et al*., 2025). Thus, the CCSA vs. L (or L^2^) trade-off provides a clearer, more mechanistic representation of the strategic choice in resource acquisition morphology.

The formation of the “conservation axis” (RTD–RN trade-off) can be mechanistically traced to a fundamental decision on C investment at the cellular level. The stele transport capacity of a single root sets an upper limit not only on the total cortex volume (Vc), but also on the total metabolic capacity that can be sustained within that volume. This implies that the potential metabolic capacity per unit cortex volume is inherently capped. Root nitrogen has conventionally been used as a proxy for metabolic capacity (Krouk *et al*., 2010; Reich *et al*., 2008; Steinkellner *et al*., 2007; Vives-Peris *et al*., 2020). Accordingly, if the total nitrogen content represents the total maximum metabolic capacity and remains relatively constant, it follows that the nitrogen content per unit cortex volume (Nv) is also relatively constant. It is assumed here that the metabolic rate and nitrogen content of the stele are sufficiently low to be negligible. Within this fixed metabolic capacity per unit volume, the plant faces a pivotal choice: how much dry mass (primarily C-rich cell walls) to invest into constructing that unit volume of cortex? This decision is executed primarily by increasing the proportion of cell wall per unit volume, i.e., by increasing the cortical cell wall thickness of individual cells or increasing the number of cells per unit volume (Zhang et al., 2024), directly dictates the trade-off between persistence and activity: (i) Allocating more carbon to thick, lignified cell walls or increasing cell number results in a high mass of structural material per unit volume—high root tissue density (RTD). However, this dense packing leaves less volume for cytoplasm and enzymes, diluting the nitrogen content per unit dry mass and leading to low RN. This is a slow-return, high-persistence strategy. (ii) Minimizing cell wall investment yields thin-walled cells with large lumens, resulting in low RTD. The saved volume is filled with metabolically active components, concentrating nitrogen into a smaller dry mass pool and yielding high RN. This is a fast-return, high-activity strategy. Under the constraint of a fixed metabolic capacity per unit volume, RTD and RN become inversely and obligatorily linked: i.e., Nv=RN × RTD ≈ constant. This forced choice, mediated by cell wall investment, is the direct mechanistic origin of the conservation axis. This framework also accommodates the suggestion that root water concentration (RWC) may be a more direct metabolic proxy (Li *et al*., 2026), where the trade-off would be expressed as Wv (the water content per unit cortex volume)= RWC × RTD.

The independence (orthogonality) of the collaboration and conservation axes in the RES arises from the decoupling of their governing constraints and processes. The collaboration axis stems from a macro-morphological investment trade-off (CCSA vs. L) under a constrained total cortex volume, determining the *size* of the metabolic “factory” (cortex volume). The conservation axis stems from a cellular construction trade-off (cell wall vs. cytoplasmic investment) under a fixed metabolic capacity per unit volume, determining the *operating mode* (durable vs. fast) of that factory. This is somewhat similar to the cell wall thickness vs. cell number independence proposed by Zhang et al. (2024). This mechanistic separation allows the two strategic dimensions to vary independently, explaining the fundamental bi-dimensionality of the RES (Fig. 5).

## CONCLUSION AND FUTURE PERSPECTIVES

This study introduces a three-dimensional C–nutrient functional balance model for first-order roots by explicitly incorporating root length. Subtropical woody species closely match model predictions, but herbaceous plants from the Qinghai–Tibetan Plateau deviate, likely due to extreme environmental impacts on transport or metabolism (Zheng *et al*., 2025). Validation across different plant life forms revealed a pronounced trade-off between CCSA and root length investment. Woody species tend to adopt a “thick and short” strategy, relying more heavily on mycorrhizal symbiosis for nutrient acquisition, whereas herbaceous species exhibit a “thin and long” strategy, emphasizing rapid self-reliance soil exploration. More importantly, this framework offers a unified mechanistic explanation for the two principal axes of the RES. The collaboration axis emerges from a macro-morphological trade-off (CCSA vs. L or CCSA vs. L^2^) under the constrained total cortex volume, distinguishing “outsourcing” to mycorrhizae from independent soil exploration. The conservation axis (RTD–RN trade-off) originates from a cellular construction strategy under a fixed metabolic capacity per root volume (Nv ≈ RN × RTD); plants decide how much C to invest in cell walls: high investment yields durable, high-RTD, low-RN roots (slow strategy); low investment yields metabolically active, low-RTD, high-RN roots (fast strategy).

Due to the current scarcity of first-order root length data, the three-dimensional C – nutrient functional balance model proposed in this study cannot yet be validated over a wider range. Nevertheless, to realize the full potential of the model, two critical extensions are needed: (i) scaling the model from single roots to hierarchical root modules. We hypothesize that the transport capacity of a second-order root constrains the combined cortex volume and metabolic load of all its attached first-order roots. Similarly, a third-order root would constrain its entire downstream branch. This hierarchical scaling would formally link single-root anatomy to branch-integrated function, providing a quantitative framework to predict resource functional balance across root orders. (ii) Extending from AM-centric to a general theory encompassing EM and other types. Our model is conceptually rooted in the anatomy of arbuscular mycorrhizal (AM) symbiosis, where intracellular colonization makes cortical volume the key limiting resource balanced against stele transport. However, this framework does not directly translate to ectomycorrhizal (EM) plants. EM fungi colonize the root surface (Hartig net), and their success is more linked to root tip abundance and branching intensity (BI) than to cortical volume (Yan *et al*., 2022; Kou *et al*., 2026), explaining the weak cortex-stele allometry in EM roots (Zhang *et al*., 2023). For EM species, a parallel theory needs development, where the stele capacity of a branch may constrain the total number of root tips (or colonizable surface area) it can support, which in turn dictates the potential size and C cost of the external fungal network. Future work must reconceptualize the balance around root tip production and the economy of the extraradical mycelium. Ultimately, integrating these extensions—across root orders and across mycorrhizal types—will pave the way for a general theory of root functional design, explaining how plants assemble entire root systems to balance acquisition, transport, and symbiosis across diverse environments. Overcoming the severe scarcity of order-specific length, anatomical data and such from intact root branches is paramount for doing these.

## ACKNOWLEDGEMENTS

We are grateful to all the authors, site investigators and their funding agencies contributing to the dataset utilized in the present study. This study was supported by the National Natural Science Foundation of China (U25A20640).

## DATAACCESSIBILITY STATEMENT

All data used in this analysis are tabulated in the Supporting Information.

## SUPPORTING INFORMATION

Additional Supporting Information may be downloaded via the online version of this article at Wiley Online Library (www.ecologyletters.com).

## REFERENCES

Bartlett, M.K., Scoffoni, C. & Sack, L. (2012). The determinants of leaf turgor loss point and prediction of drought tolerance of species and biomes: a global meta-analysis. Ecology Letters, 15, 393–405.

Bergmann, J., Weigelt, A., van Der Plas, F., Laughlin, D.C., Kuyper, T.W., Guerrero-Ramirez, N.R., et al. (2020). The fungal collaboration gradient dominates the root economics space in plants. Science Advances, 6, 1–9.

Berta, G., Fusconi, A., Trotta, A. & Scannerini, S.J.N.P. (1990). Morphogenetic modifications induced by the mycorrhizal fungus *Glomus strain* E3 in the root system of *Allium porrum* L. New Phytologist, 114, 207–215.

Cao, J.J., Wang, J.J., Yang, Q.P., Guo, B.L., Colombi, T., Valverde-Barrantes, O.J., et al. (2025). Root anatomy governs bi-directional resource transfer in mycorrhizal symbiosis. Nature Communications, 16, 1–9.

Chen, G.S., Hobbie, S.E., Reich, P.B., Yang, Y.S. & Robinson, D. (2019). Allometry of fine roots in forest ecosystems. Ecology Letters, 22, 322–331.

Chen, Q., Xu, F., Ai, Q.L., Chen, J.L. (2012). Numerical Simulation of Water Transport through Vessel Perforation Plates. Applied Mechanics & Materials, 195–196, 645-650.

Colmer, T.D. & Greenway, H. (2011). Long transport in seminal and adventitious roots of cereals during O_2_ deficiency. Journal of Experimental Botany, 62, 39–57.

Colombi, T., Chakrawal, A. & Herrmann, A.M. (2022). Carbon supply-consumption balance in plant roots: effects of carbon use efficiency and root anatomical plasticity. New Phytologist, 233, 1542–1547.

Comas, L.H. & Eissenstat, D.M. (2009). Patterns in root trait variation among 25 co-existing North American forest species. New Phytologist, 182, 919–928.

Craine, J.M., Tilman, D., Wedin, D., Reich, P., Tjoelker, M. & Knops, J. (2002). Functional traits, productivity and effects on nitrogen cycling of 33 grassland species. Functional Ecology, 16, 563–574.

Eissenstat, D.M. & Achor, D.S. (1999). Anatomical characteristics of roots of citrus rootstocks that vary in specific root length. New Phytologist, 141, 309–321.

Eissenstat, D.M., Wells, C.E., Yanai, R.D. & Whitbeck, J.L. (2000). Building roots in a changing environment: implications for root longevity. New Phytologist, 147, 33–42.

Fan, A., Wang, X., Yan, X., Chen, T., Jiang, Q., Jia, L. et al. (2024). Does a whole plant conservation gradient exist within a subtropical broadleaved evergreen forest? Frontiers in Plant Science, 15: 1481323.

Gaudinski, J.B., Torn, M.S., Riley, W.J., Dawson, T.E., Joslin, J.D. & Majdi, H. (2010). Measuring and modeling the spectrum of fine-root turnover times in three forests using isotopes, minirhizotrons, and the Radix model. Global Biogeochemical Cycles, 24, 1–17.

Guo, X.Y., Luo, J.P., Du, Y.L., Li, J.X., Liu, Y.K., Liang, Y.C., et al. (2021). Coordination between root cell wall thickening and pectin modification is involved in cadmium accumulation in *Sedum alfredii*. Environmental Pollution, 268, 1–11.

Hacke, U.G., Sperry, J.S. & Pittermann, J. (2004). Analysis of circular bordered pit function - II. Gymnosperm tracheids with torus-margo pit membranes. American Journal of Botany, 91, 386–400.

Huo, C.F. & Cheng, W.X. (2019). Improved root turnover assessment using field scanning rhizotrons with branch order analysis. Ecosphere, 10, 1–11.

Jackson, R.B. & Caldwell, M.M.J.E. (1993). The scale of nutrient heterogeneity around individual plants and its quantification with geostatistics. Ecology, 74(2), 612–614.

Kong, D.L., Ma, C.G., Zhang, Q., Li, L., Chen, X.Y., Zeng, H., et al. (2014). Leading dimensions in absorptive root trait variation across 96 subtropical forest species. New Phytologist, 203, 863–872.

Kong, D.L., Wang, J.J., Valverde-Barrantes, O.J. & Kardol, P. (2021). A framework to assess the carbon supply-consumption balance in plant roots. New Phytologist, 229, 659–664.

Kong, D.L., Wang, J.J., Wu, H.F., Valverde-Barrantes, O.J., Wang, R.L., Zeng, H., et al. (2019). Nonlinearity of root trait relationships and the root economics spectrum. Nature Communications, 10, 1–9.

Kong, D.L., Wang, J.J., Zeng, H., Liu, M.Z., Miao, Y., Wu, H.F., et al. (2017). The nutrient absorption-transportation hypothesis: optimizing structural traits in absorptive roots. New Phytologist, 213, 1569–1572.

Kotula, L., Clode, P.L., Striker, G.G., Pedersen, O., Läuchli, A., Shabala, S., et al. (2015). Oxygen deficiency and salinity affect cell-specific ion concentrations in adventitious roots of barley (Hordeum vulgare). New Phytologist, 208, 1114–1125.

Kou, L., Zuo, W., Freschet, G.T., Zheng, J., Ma, N., Lambers, H., et al. (2026). Toward refining and contextualizing the root economics space. Trends in Ecology & Evolution, Advance online publication. 10.1016/j.tree.2026.02.003.

Krouk, G., Crawford, N.M., Coruzzi, G.M. & Tsay, Y.F. (2010). Nitrate signaling: adaptation to fluctuating environments. Current Opinion in Plant Biology, 13, 265–272.

Lambers, H., & Oliveira, R.S. (2019). Plant Physiological Ecology (3rd ed.). Springer Nature Switzerland AG, Cham, 1–721.

Li, F.L., Hu, H., McCormlack, M.L., Feng, D.F., Liu, X. & Bao, W.K. (2019). Community-level economics spectrum of fine-roots driven by nutrient limitations in subalpine forests. Journal of Ecology, 107, 1238–1249.

Li, H., Carmona, C.P., Niu, S.L., Wright, I.J., Zhou, Y., Peñuelas, J., et al. (2026). The overlooked role of root water content in the root economics space. Nature Plants, 12, 1–19.

Li, H.B., Liu, B.T., McCormack, M.L., Ma, Z.Q. & Guo, D.L. (2017). Diverse belowground resource strategies underlie plant species coexistence and spatial distribution in three grasslands along a precipitation gradient. New Phytologist, 216, 1140–1150.

Ma, Z.Q., Guo, D.L., Xu, X.L., Lu, M.Z., Bardgett, R.D., Eissenstat, D.M., et al. (2018). Evolutionary history resolves global organization of root functional traits. Nature, 555, 94–97.

Matthus, E., Zwetsloot, M., Delory, B.M., Hennecke, J., Andraczek, K., Henning, T. et al. (2025). Revisiting the root economics space—its applications, extensions and nuances advance our understanding of fine-root functioning. Plant and Soil, 514, 1–27.

Meyers, B., Friaud, R., Kassie, D., Scherer-Lorenzen, M. & Freschet, G.T. (2025). Soil nutrient availability rather than spatial nutrient heterogeneity shapes the intraspecific response of root architectural, morphological and mycorrhizal traits in *Vaccinium myrtillus*. Journal of Ecology, 113, 3678–3699.

Poot, P. & Lambers, H. (2003). Are trade-offs in allocation pattern and root morphology related to species abundance? A congeneric comparison between rare and common species in the south-western Australian flora. Journal of Ecology, 91, 58–67.

Pregitzer, K.S., DeForest, J.L., Burton, A.J., Allen, M.F., Ruess, R.W. & Hendrick, R.L. (2002). Fine root architecture of nine North American trees. Ecological Monographs, 72, 293–309.

Preston, K.A., Cornwell, W.K. & DeNoyer, J.L. (2006). Wood density and vessel traits as distinct correlates of ecological strategy in 51 California coast range angiosperms. New Phytologist, 170, 807–818.

Reich, P.B., Tjoelker, M.G., Pregitzer, K.S., Wright, I.J., Oleksyn, J. & Machado, J.L. (2008). Scaling of respiration to nitrogen in leaves, stems and roots of higher land plants. Ecology Letters, 11, 793–801.

Roth, A. (1996). Water transport in xylem conduits with ring thickenings. Plant Cell & Environment, 19(5), 622–629.

Schneider, H.M., Strock, C.F., Hanlon, M.T., Vanhees, D.J., Perkins, A.C., Ajmera, I.B., et al. (2021). Multiseriate cortical sclerenchyma enhance root penetration in compacted soils. Proceedings of the National Academy of Sciences of the United States of America, 118, 1–11.

Shen, W., Xiao, Z., Xie, Z., Qin, C., Zhu, X., Zhang, T., et al. (2025). Aberrant carbohydrate partitioning 1 modulates sucrose allocation by regulating cell wall formation in rice. Plant Journal, 123(4), 1–14.

Sperry, J.S. & Hacke, U.G. (2004). Analysis of circular bordered pit function - I. Angiosperm vessels with homogenous pit membranes. American Journal of Botany, 91, 369–385.

Sperry, J.S., Meinzer, F.C. & McCulloh, K.A. (2008). Safety and efficiency conflicts in hydraulic architecture: scaling from tissues to trees. Plant Cell and Environment, 31, 632–645.

Steinkellner, S., Lendzemo, V., Langer, I., Schweiger, P., Khaosaad, T., Toussaint, J.P., et al. (2007). Flavonoids and strigolactones in root exudates as signals in symbiotic and pathogenic plant-fungus interactions. Molecules, 12, 1290–1306.

Steudle, E. & Peterson, C.A. (1998). How does water get through roots? Journal of Experimental Botany, 49, 775–788.

Sun, Y. (2007). Effect of mycorrhiza and fertilizer on morphology of the first order roots in fraxinus mandshurica and larix gmelinii plantation. (Master’s thesis). Northeast Forestry University, Harbin, China.

Vives-Peris, V., de Ollas, C., Gómez-Cadenas, A. & Pérez-Clemente, R.M. (2020). Root exudates: from plant to rhizosphere and beyond. Plant Cell Reports, 39, 3–17.

Wahl, S. & Ryser, P. (2000). Root tissue structure is linked to ecological strategies of grasses. New Phytologist, 148, 459–471.

Wang, H.F., Wang, Z.Q. & Dong, X.Y. (2019). Anatomical structures of fine roots of 91 vascular plant species from four groups in a temperate forest in Northeast China. Plos One, 14, 1–17.

Wang, S., Zhu, L.X., Zhang, P., Wang, X.Y., Sun, H.C., Zhang, K., et al. (2025b). Synergistic optimisation of root hydraulic architecture enhances drought tolerance in cotton. Plant Cell and Environment, 48, 6662–6673.

Wang, S.Y., Comas, L.H., Reich, P.B., McCormack, M.L., Phillips, R.P., Gu, J.C., et al. (2025a). Variation of root resource acquisition and conservation strategies in a temperate forest is linked with plant growth forms. Tree Physiology, 45, 1–10.

Wang, X., Liu, X.R., Chen, S., Zhu, J., Yuan, Y.Q., Zhu, R., et al. (2025c). Elevational variation in anatomical traits of the first-order roots and their adaptation mechanisms. Plant Diversity, 47, 291–299.

Wang, Z.Q., Huang, H., Wang, H., Peñuelas, J., Sardans, J., Niinemets, Ü., et al. (2022). Leaf water content contributes to global leaf trait relationships. Nature Communications, 13, 1–9.

Wei, L.L., Xu, T.T., Ma, Z.J., Zhang, L.A., Wang, Z.Y. & Ma, F. (2024). Variation in leaf and root functional traits of Caragana jubata across different provenances in a common garden. Chinese Journal of Applied Ecology, 35, 3005–3014.

Weih, M. & Karlsson, P.S. (2001). Growth response of Mountain birch to air and soil temperature: is increasing leaf-nitrogen content an acclimation to lower air temperature? New Phytologist, 150, 147–155.

Yan, H., Freschet, G.T., Wang, H.M., Hogan, J.A., Li, S.G., Valverde-Barrantes, O.J., et al. (2022). Mycorrhizal symbiosis pathway and edaphic fertility frame root economics space among tree species. New Phytologist, 234, 1639–1653.

Zhang, S.W., He, Z., Zhang, C., Sang, T., Yan, M.F., Chen, Y.L., et al. (2025). Arbuscular mycorrhizal fungi improve maize growth and antioxidant activity by modulating phytohormone contents and metabolic pathways under drought stress. Plant Physiology and Biochemistry, 229, 1–23.

Zhang, Y., Cao, J. J., Yang, Q. P., Wu, M. Z., Zhao, Y., & Kong, D. L. (2023). The worldwide allometric relationship in anatomical structures for plant roots. Plant Diversity, 45(6), 621–629.

Zhang, Y., Cao, J.J., Lu, M.Z., Kardol, P., Wang, J.J., Fan, G.Q., et al. (2024). The origin of bi-dimensionality in plant root traits. Trends in Ecology & Evolution, 39, 1–11.

Zheng, Z., Dong, F.F., Li, Z.Y., Chen, L.J., Han, Q.W., Weemstra, M., et al. (2025). The unique root form and function on the Tibetan Plateau. New Phytologist, 248, 2280–2294.

Zheng, Z., Wang, C., Wang, Y.D., Zhang, Y.R., Valverde-Barrantes, O.J., Zhang, W.H., et al. (2024). Decoupling of uptake- and transport-related traits in absorptive roots across coexisting herbaceous species in alpine meadows. Journal of Ecology, 112, 770–783.

Zhou, M., Bai, W.M., Li, Q.M., Guo, Y.M. & Zhang, W.H. (2021). Root anatomical traits determined leaf-level physiology and responses to precipitation change of herbaceous species in a temperate steppe. New Phytologist, 229, 1481–1491.

Zhou, M., Guo, Y.M., Sheng, J., Yuan, Y.J., Zhang, W.H. & Bai, W.M. (2022). Using anatomical traits to understand root functions across root orders of herbaceous species in a temperate steppe. New Phytologist, 234, 422–434.

