## Supporting Information for "A three-dimensional carbon – nutrient functional balance model explains the formation of Root Economics Space"

### Support Information

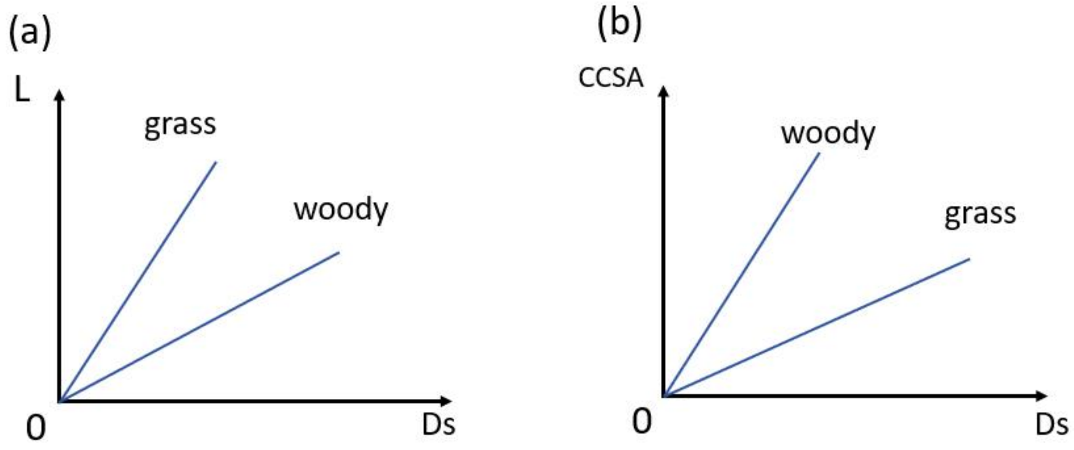


**Fig. S1** Conceptual diagram of nutrient acquisition strategies in different plants, illustrated by a comparison between woody and herbaceous life forms. (a) and (b) primarily depict the scaling relationships of root length and cortex cross-section area against stele diameter across contrasting life forms, highlighting interspecific differences in root structural investment strategies. Such variation in investment strategies may also occur within the same life form or even within a species.

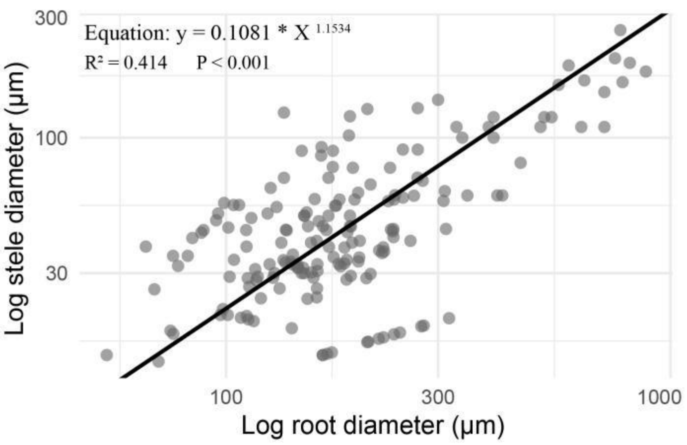


**Fig. S2** Allometric relationship between stele diameter and root diameter (stele diameter + cortex total thickness) in first-order roots from a large-scale herbaceous dataset (n = 172). Data points are shown as grey circles, and the fitted line is shown as a solid black line. The fitting equation is $y=0.1081\times x^{1.1534}$ (where y represents first-order root stele diameter, and x represents first-order root diameter (stele diameter + cortex total thickness), slope = 1.153, $R^2$=0.414, lowCI = 1.028, uppCI = 1.294.

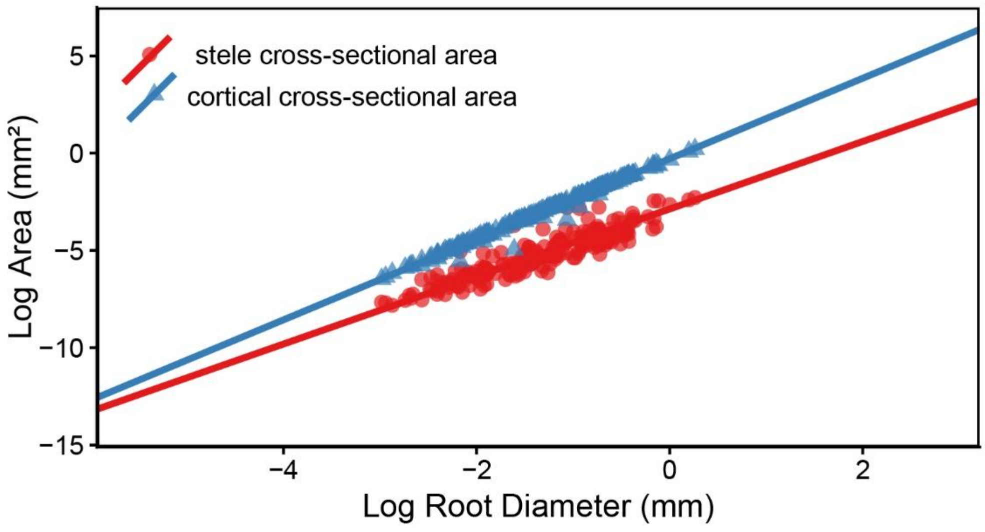


**Fig. S3** Allometric relationships between stele diameter and root diameter (stele diameter + cortex total thickness), and between cortex area and root diameter (stele diameter + cortex total thickness) in first-order roots from a large-scale woody dataset (n = 189). Data points are shown as blue circles (stele diameter vs. root diameter) and red circles (cortex area vs. root diameter). Solid lines represent fitted lines. For stele diameter vs. root diameter: slope = 1.736, $R^2$=0.829, P<0.001, lowCI = 1.635, uppCI = 1.842. For cortex area vs. root diameter: slope = 2.070, R2=0.989, P<0.001, lowCI = 2.039, uppCI = 2.102.

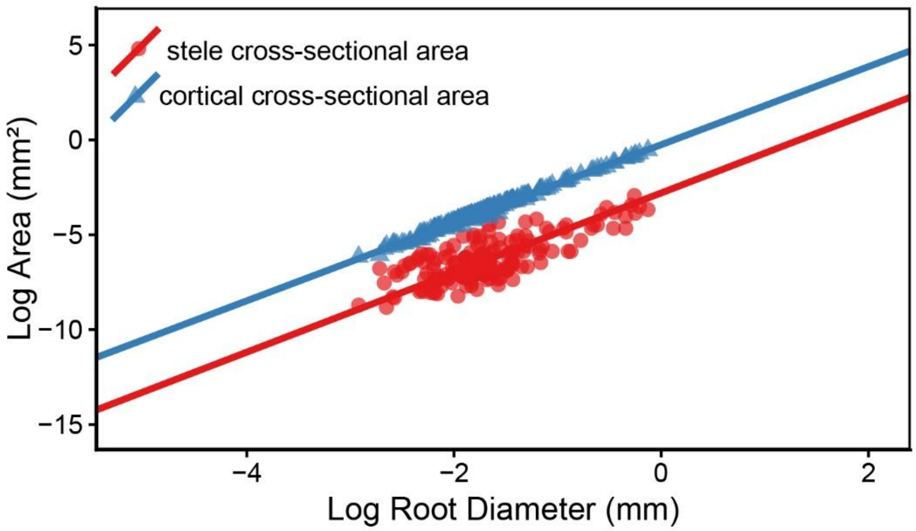


**Fig. S4** Allometric relationships between stele diameter and root diameter (stele diameter + cortex total thickness), and between cortex area and root diameter (stele diameter + cortex total thickness) in first-order roots from a large-scale herbaceous dataset (n = 172). Data points are shown as blue circles (stele diameter vs. root diameter) and red circles (cortex area vs. root diameter). Solid lines represent fitted lines. For stele diameter vs. root diameter: slope = 2.098, R2=0.576, P<0.001, lowCI = 1.902, uppCI = 2.315. For cortex area vs. root diameter: slope = 2.057, R2=0.993, P<0.001, lowCI = 2.031, uppCI = 2.084.
